# HieRFIT: Hierarchical Random Forest for Information Transfer

**DOI:** 10.1101/2020.09.16.300822

**Authors:** Yasin Kaymaz, Florian Ganglberger, Ming Tang, Francesc Fernandez-Albert, Nathan Lawless, Timothy Sackton

## Abstract

The emergence of single-cell RNA sequencing (scRNA-seq) has led to an explosion in novel methods to study biological variation among individual cells, and to classify cells into functional and biologically meaningful categories. Here, we present a new cell type projection tool, HieRFIT (**<u>Hie</u>**rarchical **<u>R</u>**andom **<u>F</u>**orest for **I**nformation **<u>T</u>**ransfer), based on hierarchical random forests. HieRFIT uses *a priori* information about cell type relationships to improve classification accuracy, taking as input a hierarchical tree structure representing the class relationships, along with the reference data. We use an ensemble approach combining multiple random forest models, organized in a hierarchical decision tree structure. We show that our hierarchical classification approach improves accuracy and reduces incorrect predictions especially for inter-dataset tasks which reflect real life applications. We use a scoring scheme that adjusts probability distributions for candidate class labels and resolves uncertainties while avoiding the assignment of cells to incorrect types by labeling cells at internal nodes of the hierarchy when necessary. Using HieRFIT, we re-analyzed publicly available scRNA-seq datasets showing its effectiveness in cell type cross-projections with inter/intra-species examples. HieRFIT is implemented as an R package and it is available at (**https://github.com/yasinkaymaz/HieRFIT/releases/tag/v1.0.0**)

## Introduction

Single-cell RNA-seq (scRNA-seq) technology has provided an unparalleled picture of the cell-to-cell complexity of biology in multicellular organisms. As technological improvements have allowed increasingly large studies, comprehensive cell atlas experiments have revealed unprecedented cell-to-cell heterogeneity and molecular dynamism of cell types across both human and model organisms (Cao et al., 2017, Rosenberg et al., 2018). Single-cell genomics have enabled tracing developmental lineages of early embryonic cells and building transcriptional landscapes of organogenesis at single-cell resolution, and uncovering novel rare cell populations (Cao et al., 2019, Tabula Muris et al., 2018).

As single-cell experiments grow in size and scope, the computational challenges associated with analyzing and interpreting these data are also growing. In particular, identifying cell types present in a sequenced population is critically important for enabling biological insight. Widely prevalent single cell analyses protocols incorporate unsupervised clustering methods as a key step in this process. For example, k-means, hierarchical clustering, KNN (k-nearest neighbor) or SNN (shared-nearest-neighbor) graphs, and Louvain community detection are all used in a variety of different packages, such as SC3 (Kiselev et al., 2017) and Seurat (Butler et al., 2018). Unsupervised clustering methods attempt to identify a consistent and biologically meaningful set of cell types or cell states in an experiment, usually via a projection of high dimensional data. These approaches have identified numerous novel subtypes (Aevermann et al., 2018, Plasschaert et al., 2018, Suo et al., 2018), although complexities of parameter optimization, number of available cells, and intrinsic noise of single-cell data can pose challenges (Kiselev et al., 2019, Tang et al., 2020).

However, unsupervised clustering approaches do not provide any rapid or automated way of defining cluster identities, which is often done by manually checking marker gene expression. In addition to being cumbersome, manual annotation depends on the robustness of a handful of a priori marker genes. When cell types are highly similar to each other transcriptomically, manual annotation may be prone to human error as reliable and obvious marker genes may not exist (Lähnemann et al., 2020). An alternative to unsupervised clustering is to use the rich information from larger atlas projects, and focus on information transfer to new studies (Wilbrey-Clark et al., 2020). While potentially faster and more accurate for cell type annotation than unsupervised clustering, especially for small-scale studies, integration and accurate information transfer between existing atlas datasets play a critical role. Supervised machine learning methods, using large cell atlas datasets as training data, provide a potential approach to automate information transfer for faster and accurate projections (Petegrosso et al., 2020).

A number of supervised classification methods have been developed, including singleCellNet (Tan and Cahan, 2019), ACTINN (Ma and Pellegrini, 2019), Garnett (Pliner et al., 2019), with different strengths and limitations. These methods differ in various aspects such as feature selection, for instance, singleCellNet trains its models with random forest after extracting a set of feature pairs from the reference data while ACTINN uses neural networks that automatically chooses the features. Garnett, on the other hand, relies only on a set of cell type specific marker genes as input independent of a reference dataset. Although the majority of these developed methods are flat classifiers, hierarchical classification has also been implemented in the single-cell context with CHETAH (de Kanter et al., 2019), and scClassify (Lin et al., 2019), which allowed intermediate class assignments, although their outputs provided limited insight into actual cell types.

Despite the rapid proliferation of cell-type assignment methods, a number of limitations still exist with current approaches. Many existing methods work best when the reference training data is composed of a few well-represented cell types, and when the query data contains a few or no novel types (Abdelaal et al., 2019a). However, an ideal classification should be able to handle many candidate cell classes, potentially hundreds, and not rely on a minimum input threshold of query data, as some single-cell protocols produce low-throughput data in which rare cell types are represented with only a few cells (Campbell et al., 2017). In addition, handling complex classification tasks by conventional methods usually involves either assigning to a cell type with low confidence or the best case is declaring them as ‘undetermined’. However, this approach underestimates the potentially informative biological signal which is often challenging to harvest and valuable to resolve experimental questions. Furthermore, considering cell types as discrete entities with clear boundaries is far from ideal as, in reality, many cells are in transitioning intermediate stages, which makes classification more compelling (Macaulay et al., 2016). Thus, alternative approaches that benefit from hierarchical consideration of cell types are required to eliminate these issues in the single-cell identity detection.

Here, we propose a new hierarchical classification approach, HieRFIT, which uses a hierarchical tree structure of reference cell clusters, allowing custom defined intermediate classes (internal nodes) that have biological meaning. Using this hierarchical model, we both improve HieRFIT’s ability to provide accurate cell type classification, and allow cells that cannot be accurately classified to be assigned to the best supported internal node. We implemented our approach as an R package and tested against various classification tasks.

## Methods

### Constructing the cell type decision tree with class hierarchies

A key input component of HieRFIT is the cell type hierarchical tree. We define this hierarchy as a tree, *τ*, which is a subtype of directed acyclic graphs, where each cell type or cell class is represented as a node *v*, and the connections between the nodes are edges, E. Nodes can only have a single parent, but can have multiple child nodes. The nodes in the tree, *τ*, are also asymmetric (each child node cannot be a parent of its own parent), and the tree itself is transitive (each node is also a child node of its parent’s ancestral node). From this cell type tree, we can define an ancestral hierarchy. Let *A* represent a set of all ancestral nodes of a given node and *Y* represent the set of class labels of all nodes. Then, *A*_*j*_ = {*v*_*j*_, *v*_*j*+1_, *v*_*j*+2_, …, *v*_*k*_} is the set of nodes comprising the ancestral path for node *v*_*j*_ reaching up to the root node *v*_*k*_ and *Y*_*j*_ = {*y*_1_, *y*_2_, …, *y*_*n*_} is the class label set for children of node *v*_*j*_. We define the terminal nodes with no children as leaves. To define this hierarchical tree for a given reference datasets, the user can input a cell type table in a tab delimited format with each row designates a leaf cell type from the reference dataset and columns represent the intermediate cell types to be used as internal nodes (**Supplementary Table 1**). HieRFIT can also create a *de novo* tree out of cell type distances based on their averaged gene expressions using hierarchical clustering if an input tree is not provided.

### Feature selection from reference data and local classifier training

Feature selection is performed for each local classifier (internal node) separately. Let *M* be the normalized expression matrix to be used as a training data, which is composed of genes *G* and samples (cells) *X* accompanied by a set of class types *Y*. In addition to the existing class types, an ‘OutGroup’ class that represents the other cell types is also added to the set, *Y*. ‘OutGroup’ class sample size is limited to a maximum 500 cells (same as other classes, and can be altered by user) for all nodes and is formed by randomly selecting cells from classes that are not present for the local classifier. HieRFIT selects a set of separate features, *f*_*j*_ ∈ *G*, for every internal node, *v*_*j*_ ∈ *v*, using the corresponding subset data *m*_*j*_ ⊂ *M*. Genes with very limited variation across cell types, *m*_*j*_ (*σ*^2^ < 0.01), are pre-filtered in order to eliminate non-informative features. After standardizing the expression by centering at the mean and scaling by the standard deviation, HieRFIT computes eigenvectors of the data with principal component analysis using the ‘*prcomp*’ function from the R stats package. To define features, HieRFIT first selects principal components (PCs) that usefully separate class labels *Y*_*j*_, by computing a t-test on the component scores of each PC and selecting PCs with *P* < 0.05 (following Bonferroni correction). To turn these informative PCs into highly variable feature sets, HieRFIT chooses the top 2000 variables (genes) that are most correlated with their eigen vectors based on absolute component loading values (number of top genes selected can be changed by user). The number of genes selected from each component is proportional to variance explained by the PC. Further, wilcoxon rank sum test between the class labels is applied to further select (by default) 200 differentially expressed genes (based on adjusted p-values) as features to be used in local classifier training.

HieRFIT constructs a reference classifier with multiple local classifiers, one for each of the internal nodes on the hierarchical tree. Local classifiers are created using a random forest algorithm implemented with the R package Caret, with the features selected separately for each node using the procedure described above, and with 500 trees (by default). The training sample set of each local classifier, *m*_*j*_, corresponds to the cells from reference data with cell type labels matching the class labels of the node’s children, *Y*_*j*_. The array of local classifiers is stored as an S4 object (in R) in the hierarchical organization to be used for projecting cell types on a query data.

### Sigmoid calibration and noise injection

The random forest classifier produces as output a vector of votes for each class, one from each tree. However, these vote distributions are not equivalent to class probabilities, therefore, they need to be transformed with a calibration function before they can be used as probabilities. HieRFIT implements Platt scaling (Platt, 1999) for this purpose: as the final step of creating a HieRFIT model, we construct a sigmoid function on reference data with class labels using a multinomial logistic regression implemented in the ‘nnet’ R package. This sigmoid function allows the conversion of class votes as the unprocessed output of random forest classifier to class probabilities. In order to provide a certain level of flexibility against dropout events in scRNA-seq, we also implemented a noise injection step prior to generating the sigmoid function (Zur et al., 2009). Noise injection occurs by setting expression values of a subset of randomly selected feature sets of each local classifier (by default 10% of all features) to zero.

### Asymmetric entropy-based certainty measurement

In order to convert class probabilities into class assignments, allowing for the possibility that some cells cannot be accurately assigned, we implemented a certainty function per candidate class. We used asymmetric entropy measurement (Marcellin et al., 2006) in our certainty function as follows;

Let *p*_*i*_ denote probability of class *y*_*i*_ ∈ *Y*_*j*_ at the node *v*_*j*_, then, the asymmetric entropy as a measure of uncertainty is

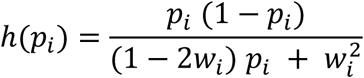

where *w*_*i*_ is probability of *y*_*i*_ at which maximum uncertainty is achieved. Note that *h* is equal to quadratic entropy of Gini when *w*_*i*_ = 0.5 in binary class modalities. From *h*, we then derived a function called ‘certainty function’, *U*, which contains an additional coefficient, *λ*, to assign directionality as follows;

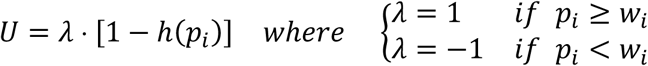

Certainty scores center at zero when *p*_*i*_ = *w*_*i*_ and range between −1 and 1 representing maximum certainties about unrelatedness and relatedness to the class, respectively. In order to obtain a set of empirical probability centroids, *W* = {*w*_1_, *w*_2_, …, *w*_*i*_, …, *w*_*n*_}, for each class, HieRFIT randomizes the feature set *f*_*j*_ of *m*_*j*_ ⊂ *M* for the corresponding node with random permutations and calculates expected probabilities of each class as the mean across iterations.

### Scoring scheme and decision rule for class projection

HieRFIT then scores each in a “top-down” manner, which refers to taking all ancestral node scores and their metrics into account beginning from root node. In order to project class labels from the HieRFIT model created using the reference dataset and a cell type tree, the first step is to obtain an array of certainty scores from all local classifiers for all class types. Let *x* ∈ *X* be a cell in a query dataset. In order to determine class type of *x*, HieRFIT generates an array of path certainty scores *U*_*ij*_(*x*) which is calculated using classification certainty scores for every node on the ancestral path, where *i* represents the class types of node *j* by traversing the tree and following the ancestral path reaching to the root *v*_*k*_ as follows;

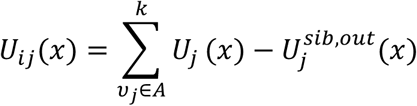

where *U*_*ij*_(*x*) is the path certainty score of cell *x* ∈ *X* for class *i* in the node *v*_*j*_, and 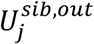 is the sum of all siblings and outgroup certainty scores for the classifier node *v*_*j*_.

During the score aggregation for decision, our rule for assigning class labels is

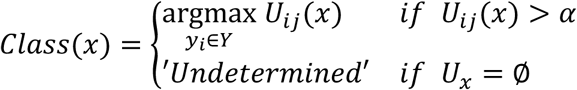

HieRFIT assigns the maximum scoring class label out of all candidate classes that pass a certainty threshold, *α*. If none of the nodes passes, “Undetermined” is returned as a class label. The certainty threshold is set to 0.05 by default, but can be changed by the user.

### Performance evaluations

For intra-dataset performance evaluation tasks, we used 5-fold cross validation, in other words, training models with 80% and testing them on 20% of data. As the evaluation metric, we calculated precision, recall, and F-measure averaged across iterations of cross validation. For inter-dataset evaluation tasks, in which training and test data originate from two separate datasets, we relied on the concordance between prior and predicted cell types of query datasets. We excluded intermediate cell type, ‘undetermined’, and multi-class assignments in metric calculations. In addition, given that HieRFIT uses a non-mandatory leaf node prediction approach and can return intermediate class labels, we also accounted for intermediate cell type assignments in performance evaluation. Therefore, we utilized precision, recall, and F-measure calculations modified for hierarchical classifications (Kiritchenko et al., 2005). For these, let *A*_*y*_ be a set of all ancestral labels for the predicted class label *y* and *A*_*θ*_ be the set of all ancestral labels for the true class *θ*_*x*_ of test sample *x*, then hierarchical precision (*hP*) and recall (*hR*) are

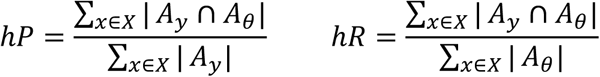

where |*A*_*v*_ ∩ *A*_*θ*_| is the number of intersecting nodes between ancestors of predicted and true class labels. Then, the hierarchical F-measure (*hF*) is

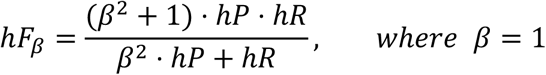

### Datasets

We analyzed PBMC scRNA-seq data from 10X Genomics with 2,700 single-cells by following standard processing workflow as instructed on Seurat online tutorials (https://satijalab.org/seurat/v3.1/pbmc3k_tutorial.html) (Butler et al., 2018). The main steps of this analysis were data quality control and normalization, identifying variable genes, data scaling, dimension reduction, clustering, finding differentially expressed genes, and assigning cell type identities to clusters based on known cell markers. We used another public PBMC scRNA-seq dataset with 68K cells (Zheng 68K) as one of the reference datasets to generate a HieRFIT model (Zheng et al., 2017b). We followed the same analysis steps in the publication (and code in GitHub https://github.com/10XGenomics/single-cell-3prime-paper). We relabeled the cell types for easier interpretation.

To evaluate the prediction performances with various data types, we selected several published single-cell datasets with available class types and expression data (**Supplemental Table 2**). We used the same cross validation folds of these datasets previously generated for benchmarking and performance evaluations through intra and inter-dataset challenges (Abdelaal et al., 2019b). These datasets originate from 10 separate scRNA-seq studies, some with multiple levels of cell type annotations.

## Results

### Overview of the algorithm

We approached the cell type classification task as a hierarchical decision problem with a set of predefined class relations. Our assumption of relationship between sub-classes and upper level classes does not necessarily have to reflect biologically defined developmental trajectories but rather represents organization of broader categories for cell type identities. HieRFIT uses multiple local random forest classifiers, organized in a higher-level hierarchical decision tree, to split the complex tasks into smaller and simpler ones. Here, we give a brief overview of the method, which is described more in details in the Methods. Given a reference expression matrix, HieRFIT first extracts the most informative principle components (PCs), which distinguish reference class types. Then, it selects a set of genes from those components as predictors based on their correlations with eigenvectors. Using the predictor set, it trains one classifier with corresponding subset data for each parent node on a user defined hierarchical tree and builds a reference classifier. In order to accurately project information from reference dataset on new experiments, we also implemented a scoring scheme for assigning class labels in a non-mandatory leaf node prediction manner, which allows us to provide intermediate cell types with broader context when data fails to provide enough resolution for more specific cell types. The uncertainty function that we derived utilizes the empirically learned background probability distribution and helps to determine whether observed probability is informative for inferring class types. Our certainty based scoring scheme properly finds the most likely ancestral path on the hierarchical cell type tree, which also provides additional confidence about identity of query cells.

### Hierarchical model construction and its algorithm

HieRFIT takes reference datasets with cell type labels along with a hierarchy of corresponding reference cell types as prior information. The hierarchical tree of cell type can be customized by the user with proper intermediate types. If the user lacks such prior information, HieRFIT can generate the hierarchy *de novo* by computing the distances between the reference cell types based on mean transcriptome expressions. To construct the reference HieRFIT model (HierMod), we implement a six step protocol that is repeated for every internal node on the hierarchical tree (**Figure 1A**). For each node a local classifier is generated using a random forest classification algorithm. To prepare the reference expression data for model training, the first step is to extract a cell data matrix that corresponds to the node. Training data is relabeled by bundling the grandchildren under the node’s children labels and adding an outgroup class as the representer of other classes. A principal component analysis using the relabeled data provides the components that are highly variable across cells and we select the components whose loadings significantly separates cells with shared type from the others. The selected significant components allow us to reduce the total number of genes to a highly variable gene set among which we select the final feature set following the Wilcoxon rank sum test. This final feature set is used in the training of the local classifier with the cell type labels (or relabels). All local classifiers are stored as an array of models which are organized in accordance with the input tree hierarchy.

**Figure 1.**
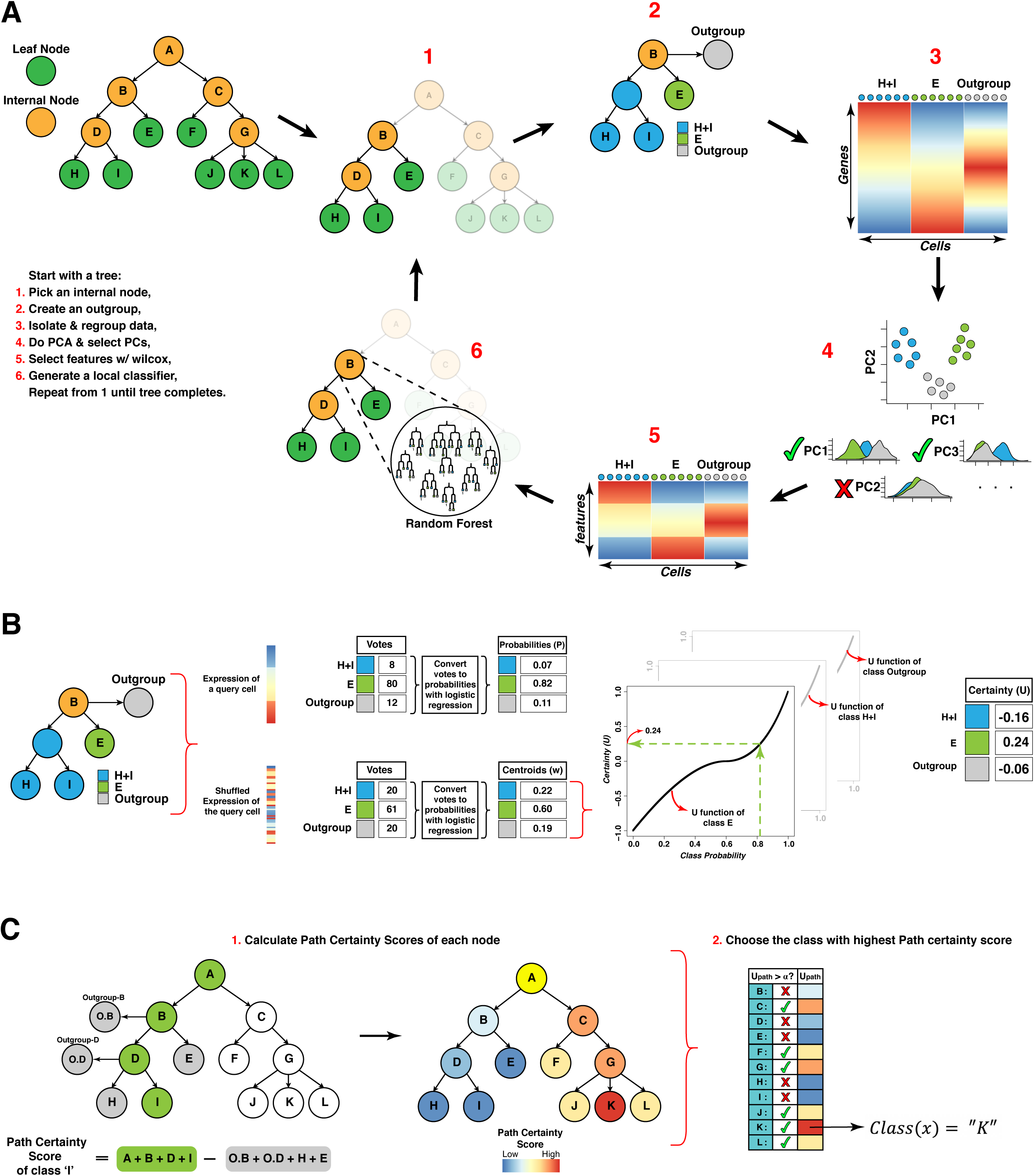
HieRFIT workflow overview. Reference model generation and query projections. Overview of HieRFIT reference model generation and prediction of a query cell class. **A)** Main process starts with obtaining a tree as a user input or creating from the data. The steps for generating the reference model with a hierarchical tree: **1**. Pick an internal node *i* (i.e. node “B”) on the tree, **2**. Re-group its children nodes and create an outgroup node for it, **3**. Extract the input expression data based on new group labels for the node, **4**. Perform Principal Component Analysis and pick the components that separate the class labels for variable feature selection, **5**. perform Wilcoxon Rank sum test to determine differentially expressed features, **6**. Train a local classifier (Random forest) with the group labels and the expression matrix with selected features. Repeat the process until all node classifiers are constructed. **B)** Query of a test cell and certainty calculations. Given an array of feature expressions of the query cell, the first step is to compute the certainty array (*U*) for the candidate classes. Votes are collected from each node *i* (i.e. node “B”) for both observed query data and its shuffled data separately. Votes are converted to probabilities using sigmoid calibration with multinomial logistic regression. Using the probability centroids (*w*_*i*_) as the outcome of the randomized array and the observed probabilities (*p*_*i*_), compute the certainty value of each class of the node (i.e the certainty of class “E” is 0.24). Repeat the process for every class of all internal nodes. **C)** Determining the cell type/class of a query cell. Step 1: Path certainty scores of each candidate class are computed using the certainty values of nodes for the given query by traversing the tree. The sum of certainty values of outgroup and sibling nodes along the path (nodes in gray) are summed and subtracted from the sum of Certainty values of nodes on the path (nodes in green). Step 2: As the final step, scores are evaluated and the maximum scoring class is returned as the outcome. If none of the classes passes the threshold, α, “Undetermined” is returned.

### Path certainty score computation and class selection

Classical measures used in decision trees such as Shannon’s entropy or quadratic entropy of Gini are not suited well for real life imbalanced data with their symmetry assumptions for equiprobability distributions among classes (Zighed et al., 2010). Therefore, we created an alternative certainty function derived from asymmetric entropy (see methods). The main stage of assigning a reference cell type to a query cell is to compute the array of certainty (U) values for each internal and terminal node (**Figure 1B**). In order to obtain such a metric, HieRFIT takes the gene expression array of the query cell and the reference model (HierMod) as inputs. Along with the gene expression array of the query cell, a randomized (shuffled 1000 times) expression array for the same query is generated. Class votes from local classifiers are obtained for both observed and shuffled expression arrays, simultaneously. Normalized class votes gathered from the local classifiers are converted to class probabilities with logistic regression (aka sigmoid calibration) function. The observed expression array is used to acquire adjusted class probabilities from each local classifier, while the shuffled expression array is used to determine class centroids. These centroids are used as certainties of the classes for the query with the observed class probabilities using the certainty function (see Methods for details).

After computing the certainty values of each node for the query, we calculate the path certainty scores for all candidate nodes by adding up the certainty values along the ancestral path and subtracting all outgroup and sibling nodes certainties (**Figure 1C**). This path score defines the final value for the nodes to be considered as candidate classes. The decision stage simply consisted of choosing the maximum scoring node among the ones whose score exceeds a certain threshold (alpha). In cases where no class exceeds the threshold, HieRFIT returns an “Undetermined”. This decision scheme permits internal nodes to be cell types of the query as well as leaf nodes that are constrained with input the reference data in the first place.

### PBMC cell type classification with HieRFIT

For demonstration purposes, we generated a reference model using the 68K PBMC single-cell dataset from Zheng et al (Zheng et al., 2017b). We created an example hierarchical tree which organized the reference cell types into two main groups, myeloid lineage and lymphoid lineage, along with the hematopoietic stem cells (HSC) using the input tree file in **Supplemental Table 1** (**Figure 2A**). Two main groups branched into further intermediate groups and general cell types. The terminal nodes comprise reference cell types from the PBMC data. We used this custom made hierarchical tree to create a HieRFIT model with maximum 500 cells per cell type in the training process.

**Figure 2.**
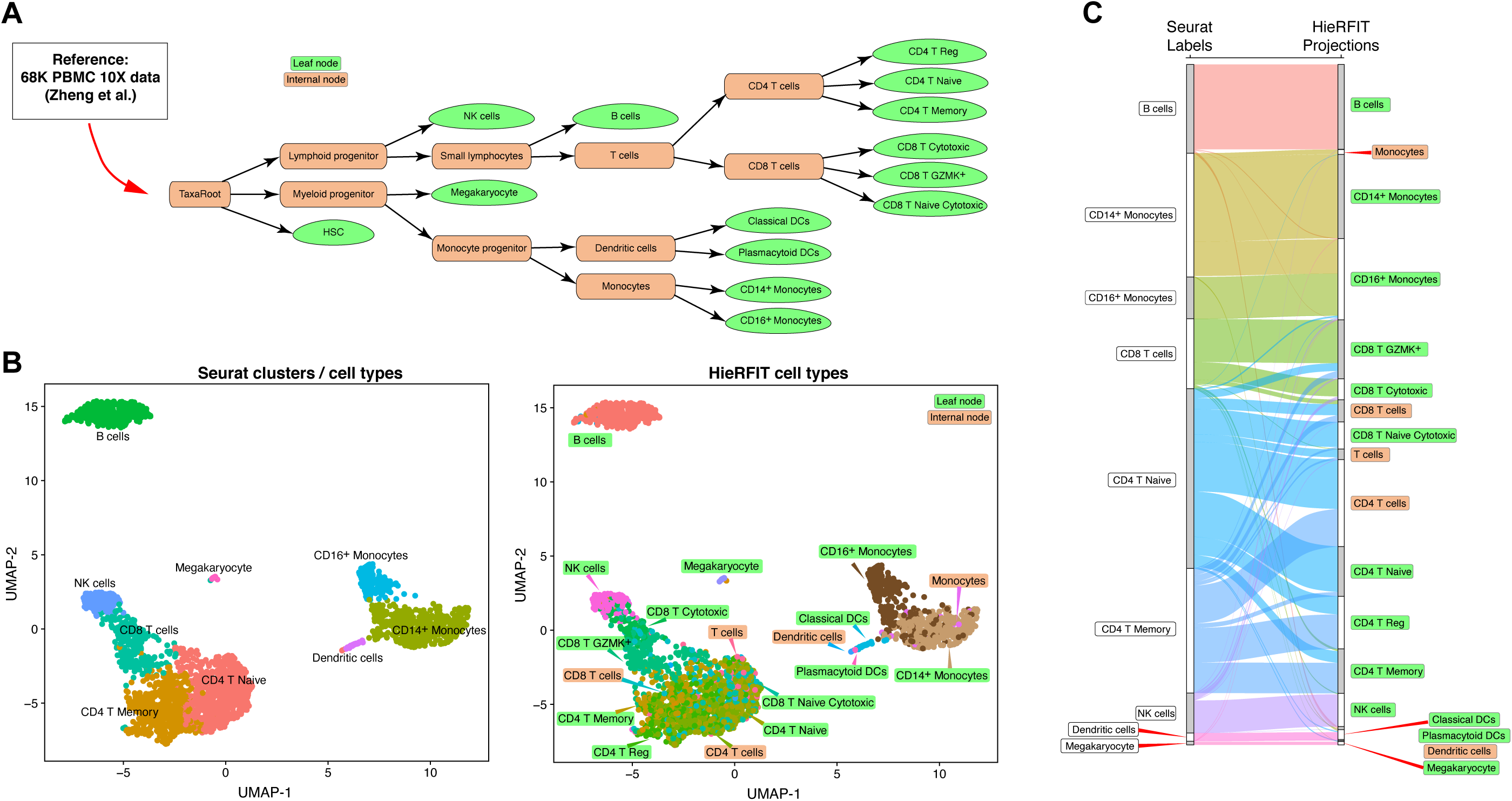
Demonstration of HieRFIT usage on a PBMC dataset. **A)** The cell type tree used in HieRFIT reference model with 68K-PBMC data. **B)** The UMAP representation of 3K-PBMC data from 10X Genomics. Cells are colored with cell types which were identified through Seurat clustering and marker expressions (left), cells are colored with HieRFIT reference cell types along with intermediate types specified in the tree file (right). **C)** Alluvial diagram demonstrating the cross comparisons of HieRFIT projections with the Seurat cell type labels. Each line connecting the two vertical black columns (left bar: prior labels, right bar: projections) represent a cell and are colored based on its HieRFIT projection type. Annotations with less than 1%, ‘HSC’ and ‘Monocyte progenitor’ were not shown.

To test our hypothesis that our hierarchical classification approach provides more accurate and meaningful results as compared to conventional way of cell type identification, we used a toy dataset, another 3K PBMC, as the query (10X Genomics, https://support.10xgenomics.com/single-cell-gene-expression/datasets/1.1.0/pbmc3k). This publicly available dataset has been generated by 10X Genomics and processed through the Seurat pipeline. We followed the same guideline as in the Seurat tutorial to identify cell types with no prior information. This manual type annotation involves several commonly accepted processes, such as finding the variable genes, dimension reduction, and clustering of cells. Finally, determining the cell types involves manually checking the marker gene expression that are differentially expressed in clusters against the rest of the groups. This process results annotation of 3K PBMC data with cell types: ‘B cells’, ‘Megakaryocytes’, ‘CD14^+^ Monocytes,’ ‘CD16^+^ Monocytes’, ‘Dendritic cells’, ‘NK cells’, ‘CD8 T cells’, ‘CD4 T Naïve’ and ‘Memory cells’ (**Figure 2B**, left). Then, we tested the hierMod we created with the 68K PBMC data by projecting the reference cell types on the same 3K PBMC dataset. HieRFIT projections labeled the cell in the query with leaf node labels as well as intermediate cell type defined in the model hierarchical tree above (**Figure 2B**, right). HieRFIT projections demonstrated a significant concordance with Seurat cell types for the distinct cell types, such as ‘B cells’, ‘Megakaryocytes’, ‘NK cells’, and ‘Dendritic cells’ (although these cells’ HieRFIT projections were “Classical DCs” rather than the parent node “Dendritic cells” on the tree). Seurat ‘CD8 T cells’ were labeled extensively with subtypes “CD8 T GZMK^+^” and “CD8 T Cytotoxic” as well as their parent node “CD8 T cells”, partially (**Figure 2C**). “CD4 T Memory” cells were labeled mainly as “CD4 T Memory” and “CD4 T Reg” in addition to the parent node “CD4 T cells”. “CD4 T Naive” cell group, on the other hand, received labels from almost all CD4 T sub-levels cell types and intermediate types such as “CD4 T cells” or even “T cells” as higher nodes on the tree. Interestingly, a group of cells within naïve CD4 T cells received CD8 T cell labels, especially “CD8 T Naïve Cytotoxic”. Similarly, a subgroup of “CD14^+^ Monocytes” were labeled as “CD16^+^ Monocytes”, while all “CD16^+^ Monocytes” were correctly labeled by HieRFIT. A small group of cells from the CD14^+^ cells were labeled as a parent node “Monocytes”.

### HieRFIT classifications are concordant with marker gene expressions

To investigate the discordance between Seurat annotations and HieRFIT projections for some of the cell groups, we further explored the marker gene expressions and their distribution. As the heatmap of the confusion matrix demonstrates, 7 out of 9 cell types were labeled with cell types by HieRFIT with more than 80% concordance (**Figure 3A**). Two of the cell types, “CD14+ Monocytes” and “CD4 T Naïve” received classification labels that resulted in 67.7% and 73.5% concordance, respectively. For the “CD4 T Naïve” cell types, we examined the expression distribution of the CD8 T cell markers, CD8A and CD8B, as well as major CD4 markers for naïve cells, IL7R and CCD7 (**Figure 3B**).

**Figure 3.**
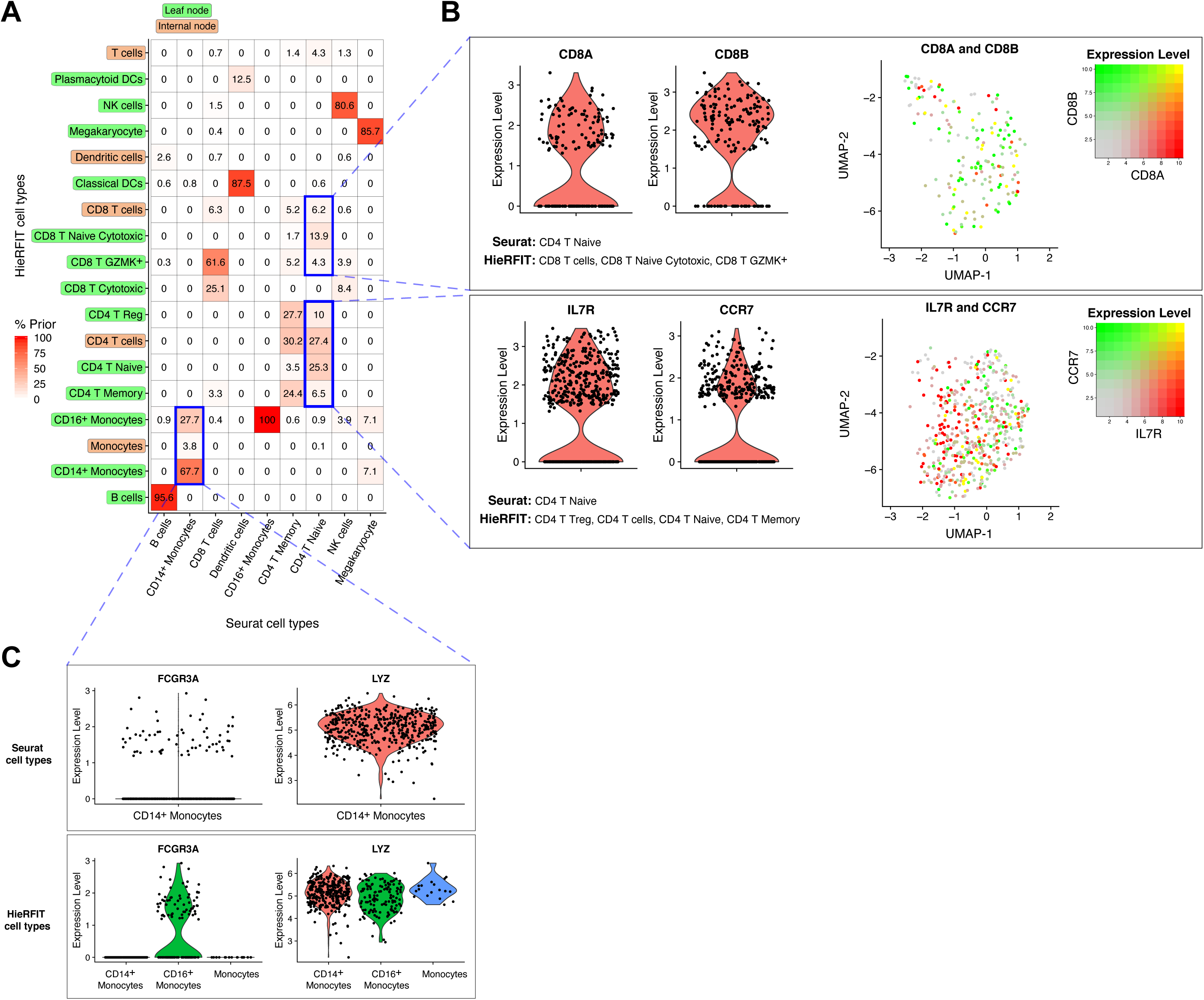
Concordance analysis of Seurat and HieRFIT classifications with gene expression of cells. **A)** The heatmap representation of the confusion matrix that summarized the projection results of the 3K-PBMC query data with percent distribution among the tree node labels. **B)** Violin plots of CD8A and CD8B genes and their co-expression values projected on UMAP representation of cells classified as “Naïve CD4 T cells” by Seurat while HieRFIT predicts them as CD8 cells or its subtypes (upper panel). Similarly, violin plots and co-expression values of IL7R and CCR7 genes projected on UMAP representation of cells predicted as CD4 T cells or its subtypes by HieRFIT in concordance with Seurat (lower panel). **C)** Normalized expression distribution of three marker genes, “LYZ” and “FCGR3A (CD16)”, markers of “monocytes” and subset “CD16 monocytes”, respectively, among the cells classified as “CD14+ Monocytes” by Seurat (upper panel). Similar violin plots for expression distribution of the same set of cells grouped based on HieRFIT projections (lower panel).

Within this group, the subset of cells which are predicted by HieRFIT to be CD8 T cells or its subtypes expressed CD8A and CD8B at high levels, suggesting these cells are properly assigned to the CD8 subtype (**Figure 3B**, upper panel - violin plots). The co-expression of the two markers also clearly showed that the significant majority of these cells in fact expressed at least one of these markers or both at the same time (**Figure 3B**, upper panel - UMAP panel with co-expression projections). This observation supports the accuracy of HieRFIT projections that, in fact, these cells are a class of CD8 T cells rather than CD4 T cells. On the other hand, the group of cells that were concordantly labeled as CD4 T cells or its subtypes carried the proper CD4 T naïve marker expressions, IL7R and CCR7, in line with their projected cell types (**Figure 3B**, lower panel – violin plots and UMAP projections).

We also investigated another group of cells with discordant predictions in “CD14+ Monocytes”. Of these monocytes, 27.7% were predicted as “CD16+ Monocytes”. When we examined the marker gene expression in cells classified as “CD14+ Monocytes” by Seurat, we observed that a significant portion of them expressed CD16 (FCGR3A) monocyte marker at high levels (**Figure 3C**, upper panel). On the other hand, HieRFIT classification of these cells demonstrated a clearer separation between these two highly similar subtypes of monocytes while preserving the major monocyte marker expression in all cells even in cells predicted with the label of the parent node, “Monocytes” (**Figure 3C**, lower panel).

### Comparative performance evaluation with intra-dataset tests

We evaluated the performance of HieRFIT on a large number of different datasets, with varying complexity, technology, and size. These include human and mouse pancreas datasets (Baron et al., 2016, Muraro et al., 2016, Segerstolpe et al., 2016, Xin et al., 2016), human PBMC (Zheng et al., 2017a), human lung cancer cell lines (Tian et al., 2019), mouse cortex and nervous system (Tasic et al., 2018, Zeisel et al., 2018) as well as whole mouse datasets from Tabula Muris consortium (2018) (**Supplemental Table 2**). We also compared its performance against other cell type classification that use supervised machine learning approaches to create a predictive model based on the training data. The first benchmarking was based on intra-dataset evaluations with 5-fold cross validation. Some of the datasets with multi-level cell type annotations were treated separately as different datasets. We calculated the mean-F1 score of each classification tool as the overall performance averaged across each cell class in the datasets. To obtain a fair benchmarking, we included only leaf node predictions of HieRFIT and excluded the intermediate node classifications in the F1-score computations.

We compared the performance of HieRFIT against 21 classification approaches with various modes using 17 unique tools. Based on the median value of the mean-F1 scores from test datasets, HieRFIT demonstrates better performance than 16 of them (**Supplemental Figure 1**). LDA, ACTINN, SingleR, SVM, singleCellNet, and SVM with rejection demonstrate comparable performance against HieRFIT (**Figure 4A**, heatmap). Both SVM and SVM with rejection option perform better HieRFIT on most datasets except two of them. Out of 18 mean-F1 scores of datasets, ACTINN is better on 12 datasets, LDA is 11, singleCellNet 10, and singleR is better on only 3 datasets compared to HieRFIT. SingleR and singleCellNet fails to complete the tasks on the complex datasets with large number of cell types, such as Zeisel (237) and AMB (92), TM (55), and Zheng datasets. 5 out of these 6 classification approaches lack an important feature, a rejection option. HieRFIT, along with LDA (with rejection), scClassify, and CHETAH, returned low levels of ‘unlabeled’ predictions while SVM (with rejection), scmap (both ‘cell’ or ‘cluster’ modes), Cell-BLAST, and scID classifications contained high level of ‘unlabeled’ results (**Figure 4A**, boxplot). Almost all of the classification tools perform the worst on Zheng PBMC (11 cell types) dataset, likely due to its intrinsic complexity.

**Figure 4.**
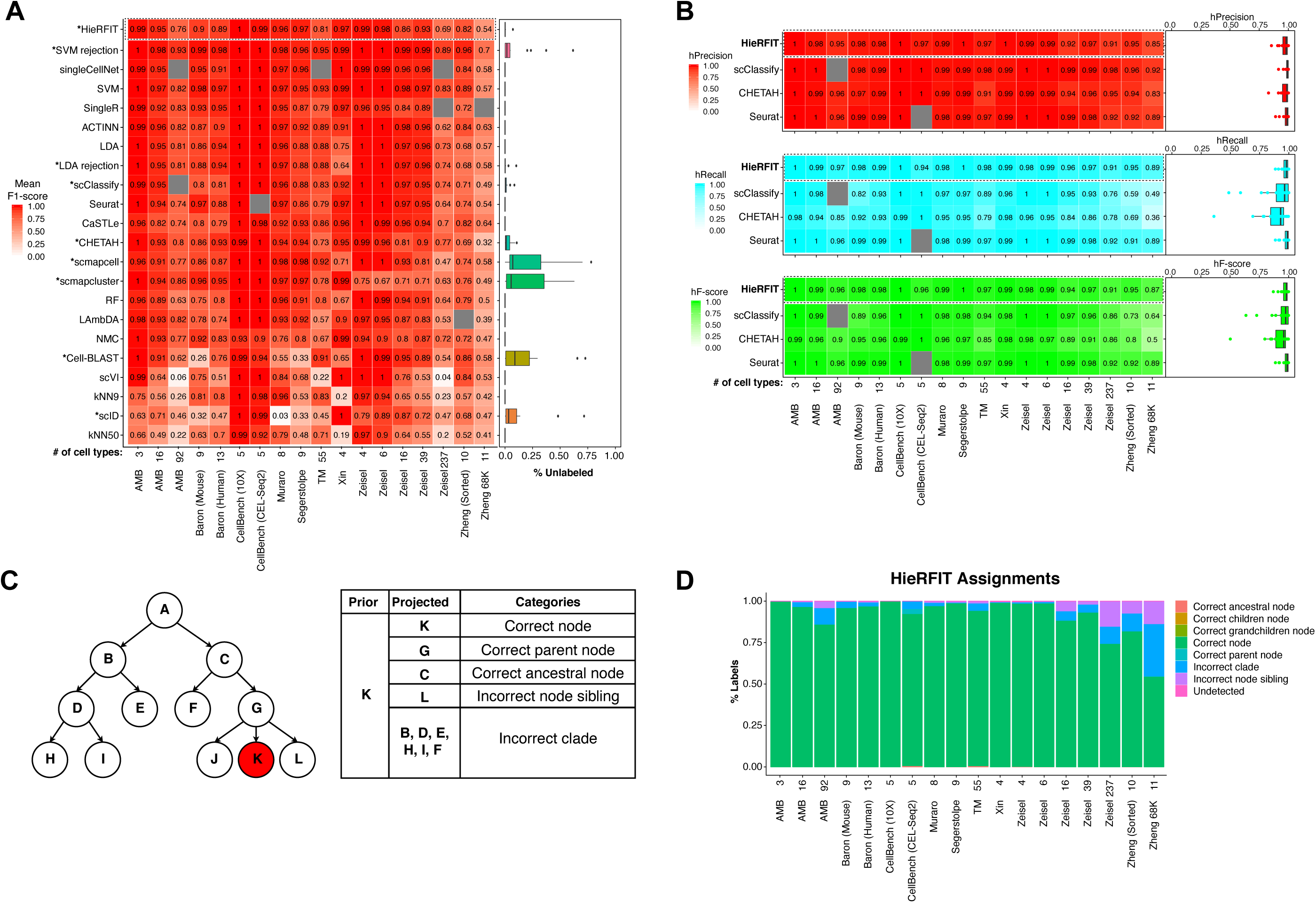
Performance on various types of datasets. Robustness against batch biases. Performance results on various types of datasets and comparative benchmarking against other cell type classification tools. **A)** A heatmap representing the mean F1 scores of each test dataset for the classification tools. The number of cell types of each dataset is shown below the columns. Failed tests without a score are grayed-out. Percent unlabeled data distribution from each test data is shown with an adjacent box plot for each tool. Asterisk (*): Classification tools with rejection option. **B)** Hierarchical precision (red), recall (cyan), and F-score (green) metrics for the tools HieRFIT, scClassify, CHETAH, and Seurat. **C)** Various categories of projected cell types by HieRFIT based on their position on tree relative to the prior label. In addition to categories in the table, “Correct children” or “Correct grandchildren” categories are also possible in case of a correct sub-level type assignment. Bar plot summarizes the distribution of these categories for HieRFIT outputs among all test datasets above. **D)** Stacked-bar plot summarizes the distribution of projection categories for HieRFIT outputs among all test datasets above.

We further explored the performance of HieRFIT in depth by comparing it to other two tools, scClassify and CHETAH, with similar hierarchical classification approaches to ours and with the most commonly used software, Seurat. We computed the hierarchical precision, recall, and F-1 score, which takes the intermediate cell type predictions into account when computing the performance metric. To be fair to the other tools, we used the same hierarchical tree that HieRFIT used in the computation of the hierarchical metrics for the other tools. We obtained the results for hierarchical precision, recall, and F-1 score measurements from the 18 intra-datasets through 5-fold cross-validations.

HieRFIT and the other three classification tools demonstrate high levels of hierarchical precision in all datasets, >91%, except ‘Zheng’ datasets (**Figure 4B**, upper panel). However, scClassify fails to return the results for AMB (92) and Seurat fails to identify enough significant anchors for “CellBench (CEL-Seq2)” dataset. On the other hand, HieRFIT, returns class predictions with consistently high recall rates (> 89%) for all datasets while scClassify and CHETAH showed significantly lower recalls especially on tasks with complex datasets with large number of cell types (**Figure 4B**, middle panel). As the performance metric that takes precision and recall into account, hierarchical F-1 score clearly demonstrates that HieRFIT performs at consistent levels and comparable to Seurat classifications (**Figure 4B**, lower panel).

To better evaluate the HieRFIT results in the hierarchical classification context, we categorized the projected cell types based on their positions on the reference tree (**Figure 4C**). These categories reflect the level of prediction accuracy relative to the hierarchical relationship defined as cell type similarities. These categories are as follows: The projection cell type is categorized as ‘Correct node’ if it is same as the true cell type (prior), as ‘Correct parent’ if it is parent of true cell type, as ‘Correct ancestral node’ if it is on the ancestral path (excluding parent node) of true type, as ‘Incorrect sibling’ if it is a sibling of true type, and as ‘Incorrect clade’ is if it is any other node with an unshared parent as true label. Using these schemes, we checked the distributions of categorized HieRFIT projections for each intra-dataset task (**Figure 4D**). HieRFIT returns a large proportion of correct leaf nodes for the majority of datasets. Even for the complex datasets, such as TM (55), AMB (92), and Zeisel (237), the correct leaf node rates are 95%, 85%, and 75%, respectively, while Zheng PBMC dataset (11 cell type) results in inferior profile due it its complexity with high rates of incorrect sibling and clade assignments. The rates of assignments from other categories are relatively lesser simply due to the intra-dataset tasks using part of the same data to test the performance.

### Robustness against various scRNA-seq methods (PBMC bench)

To evaluate the performance of HieRFIT on inter-dataset tasks in which the reference model is built on a dataset completely different from the test set, we utilized another public data collection generated for PBMC from two individuals (Ding et al., 2019). This type of inter-dataset tasks provide more realistic results as they reflect real life usage better. PBMC1 and PBMC2 samples were split into multiple subsets and sequenced with 8 different versions of single-cell RNA-seq methods. In the experiment, we trained the model using data from one method and tested it on datasets generated using other methods (and on the second of the sample pairs in case same methods). As hierarchical precision, recall, and F1-score distributions on all of the combinations, HieRFIT performs consistently well on almost all tasks with average 85% rates (**Figure 5**). On the other hand, Seurat and CHETAH, sacrifices extensive precision and recall rates, respectively, on many tests while scClassify performs slightly better than those two. However, HieRFIT outperforms all with the highest hF1-score in average across multiple tasks. Although HieRFIT’s worst performance appears to be the model generated with inDrop data when tested on CEL-Seq data as 64% hF-score, it is still comparably better than its contenders. Overall, these results show that HieRFIT exhibits robustness against various batch effects due to different scRNA-seq methods and performs better than other tools in various aspects.

**Figure 5.**
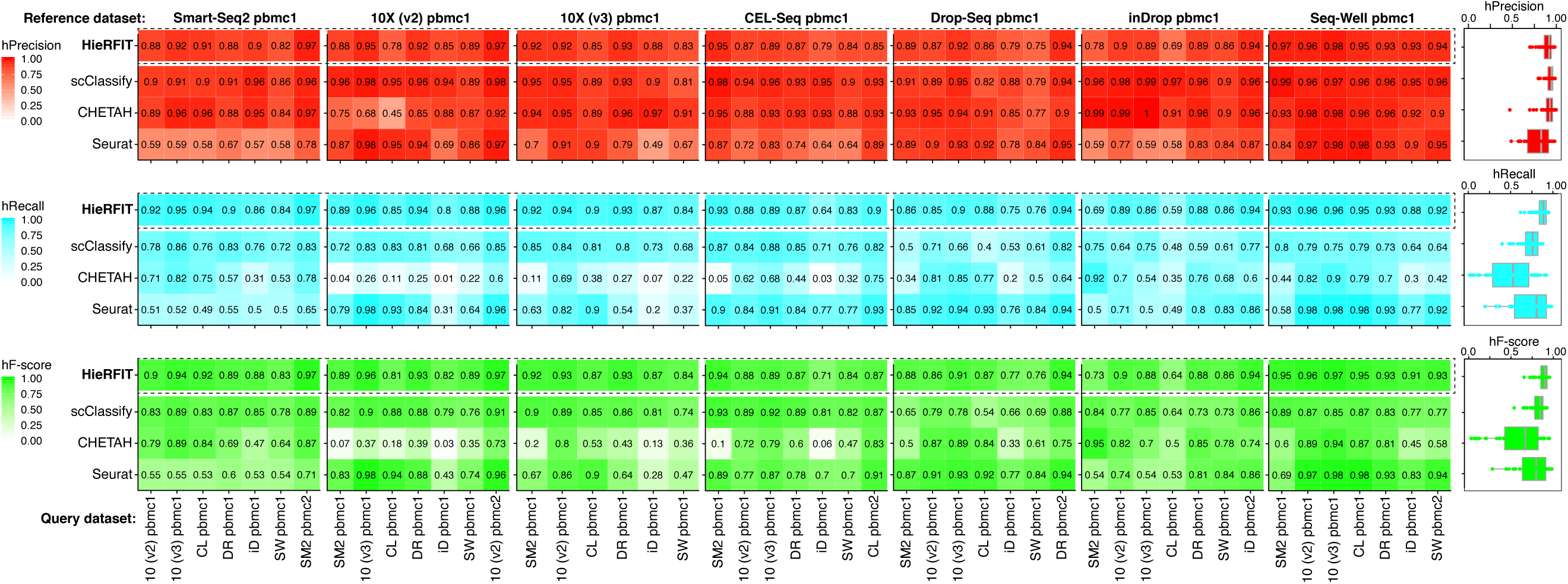
Robustness against batch biases. Hierarchical precision, recall, and F1-score values of HieRFIT and 3 tools for comparing the performances in various batches with inter-dataset tests. At each iteration, a dataset from paired PBMCs produced with a scRNA method was used to generate the reference model and tested on the second pair of the PBMCs.

## Discussion

Defining cell types is a fundamental and complex challenge in single-cell biology, which is becoming increasingly difficult as the diversity of single-cell experiments increases. In this study, we attempted to address one of the challenges in the developing field of scRNA-seq with an alternative perspective. Hierarchical classification, as opposed to common flat classifiers, is currently generating more interest in the community because it takes the cell type relationships into account in addition to providing more insight into intermediate cell types (Wu and Wu, 2020). The hierarchical approach has been used in many other fields including medical sciences (Dimitrovski et al., 2011).

In this work, we hypothesized that hierarchical organization of cell types and class relationships will provide more accurate decisions compared to flat classification approaches. We, then, implemented our approach as a user-friendly R package and evaluated its performance with commonly used public datasets. We demonstrated HieRFIT’s better classification of PBMC cell types, even in low abundances, in concordance with their marker gene expression profiles as opposed to manual annotations by widely used software, Seurat. In addition, the performance evaluations against other available single-cell classification tools and machine learning algorithms showed that HieRFIT provided the most reasonably accurate results. HieRFIT’s performance stayed stable across various types of datasets produced with different methods while other tools sometimes showed diminished accuracy, in particular, inter-dataset tasks. With its ‘divide and conquer’ approach, HieRFIT was able handle very complex tasks with a large number of cell types and total cells without any issue.

Furthermore, HieRFIT showed consistently better performance on classification challenges against two other tools, CHETAH and scClassify, which have similar hierarchical classification approaches. Both CHETAH and scClassify learn tree topologies directly from reference data as CHETAH builds a binary tree of cell types using average linkage distances based on their Spearman correlations while scClassify uses hierarchical ordered partitioning and collapsing hybrid (HOPACH) algorithm for tree construction which allows multi-children nodes. However, both tools define the intermediate cell types with labels that are hard to interpret. HieRFIT on the other hand covers both approaches by providing users an option, in addition to ability to create a de novo tree, to define a tree containing intermediate nodes with meaningful cell labels as opposed to other tools.

We implemented the ‘local classifier per parent node’ (LCPN) approach in HieRFIT as opposed to the global classifier approach that takes the entire tree topology into a single model. Hierarchical classification implemented in LCPN attitude has been reported to have better accuracy as compared to flat classifiers (Gauch et al., 2009, Jin et al., 2008, Xiao et al., 2007). In addition, using various combinations of different classification algorithms as local classifiers has previously been reported (Secker et al., 2007). One important feature of HieRFIT originates from its ‘non-mandatory leaf node prediction’ based decision scheme which allows intermediate nodes to be assigned as well. Our decision rule is based on choosing the best scoring node on the tree. Alternative decision approaches have been proposed, e.g. ‘sequential boolean decision rule’ which chooses child nodes, starting from root, until reaching to a leaf node (Bryant, 1992). However, this approach might be prone to error propagation more than other top-down approaches. The challenge is to properly combine local classifiers so that their unbiased outputs can be used for the decision making process. It is commonly known that machine learning based classifiers are prone to imbalanced class sizes in addition to other intrinsic biases such as batch effect. To account for these, we utilized a certainty function derived from asymmetric entropy which provided precise confidence estimations about class assignment. The path certainty metric accumulates higher scores when correct cell types are picked along the ancestral path while their siblings and out-groups behave as antagonists. Thus, accumulated confidence allows better decision regardless of the complexity of data or tree topology.

As all other computational approaches, HieRFIT also has several limitations. First of all, it requires a reference data with properly annotated cell types. Although relying on reference data and its cell types can introduce biases due to inconsistencies in annotations, HieRFIT’s ensemble based classifiers, random forest, can compensate for subtle fluctuations. Reference based classification approaches usually miss the opportunity to discover novel cell types due to their dependency on prior information. Another limitation of reliance on reference data is that some cell types are represented with low numbers of cells. However, with the fast development of new methods, single-cell based atlas projects provide exponentially increasing datasets. Secondly, HieRFIT relies on a user provided tree, a predefined class hierarchy, and assumes that the tree topology reflects biological cell type relationships with their underlying gene expression profiles in the reference data. To prevent senseless results, users must be cautious about providing a tree topology for classification purposes with HieRFIT. If a user skips to provide a cell type tree, creating a class hierarchy by learning from data (e.g. by hierarchical clustering) can also be limited since similarity driven hierarchy is prone to data specific artifacts and over-fitting.

In this study, we proposed to utilize hierarchical relationships between cell types to better harvest biological information and provide more insight about the cell type identities. HieRFIT provides stable and accurate cell type classification of single-cell RNA-seq data with hierarchical manner. It will contribute to the field not only by providing a new perspective and faster cell type projections from larger atlas projects but also allowing cross comparisons across various datasets effectively.

## Acknowledgements

Special thanks to Aaron Kitzmiller, Hansaim Lim, and Varenka Rodriguez Diblasi.

## Software availability

HieRFIT is available as an R package through GitHub (https://github.com/yasinkaymaz/HieRFIT).

## Supplemental Tables

**Supplemental Table 1:** An example tab-separated cell type table to be used as an input for tree construction and creating a reference model (referencing 68K PBMC dataset). Each row specifies all ancestral/intermediate cell types of each reference cell type (leaves at the end of rows).

**Supplemental Table 2:** Datasets used in the benchmarking analysis.

## Supplemental Figures

**Supplemental Figure 1**. Boxplot for mean F1 scores distribution of each classification tool as an outcome of 18 datasets with 5-fold cross-validation tests (upper plot). Percent unlabeled data distribution from each test data (middle plot). A heatmap showing the Mean F1 scores of each test dataset for the classification tools (lower panel). Failed tests without a score are grayed-out.

## Notes

### Competing Interest Statement

The authors have declared no competing interest.

